# Axon guidance genes are regulated by TDP-43 and RGNEF through the rate of long-intron processing

**DOI:** 10.1101/2023.12.05.570131

**Authors:** Yasmine Abbassi, Sara Cappelli, Eugenio Spagnolo, Alice Gennari, Giulia Visani, Simone Barattucci, Francesca Paron, Cristiana Stuani, Cristian A. Droppelmann, Michael J. Strong, Emanuele Buratti

**Author notes:** Correspondence to: Emanuele Buratti, Molecular Pathology Group, International Centre for Genetic Engineering and Biotechnology (ICGEB), AREA Science Park, Padriciano 99, 34149 Trieste, Italy. Abbreviations: ActD = Actinomycin D; ALS = Amyotrophic Lateral Sclerosis; CHX = cycloheximide; DEGs = differentially expressed genes, FC = Fold Change, FTD = frontotemporal dementia, FTLD = Frontotemporal Lobar Degeneration, FUS = fused in sarcoma, hnRNPs = heterogeneous nuclear ribonucleoproteins, NMD = nonsense mediated decay, qPCR = quantitative PCR, RNA-IP = RNA-Immunoprecipitation, RGNEF = Rho guanine nucleotide exchange factor, TDP-43 = TAR DNA-binding protein 43 kDa, *UBE2E3* = Ubiquitin Conjugating Enzyme E2 E3, UNC13A = Unc-13 homolog.

## Abstract

Rho guanine nucleotide exchange factor (RGNEF) is a guanine nucleotide exchange factor (GEF) mainly involved in regulating the activity of Rho-family GTPases. Previous work has shown that RGNEF inclusions in the spinal motor neurons of ALS patients co-localise with TDP-43, the major RNA binding protein aggregating in the brain and spinal cord of ALS patients. To further characterise their relationship, we have compared the transcriptomic profiles of neuronal-like cells depleted of TDP-43 and RGNEF and show that these two factors predominantly act in an antagonistic manner when regulating the expression of axon guidance genes. From a mechanistic point of view, our experiments show that the effect of these factors on the processivity of long introns can explain their mode of action. Our findings highlight that neurodegenerative processes at the RNA level can often represent the result of combinatorial interactions between different RNA binding factors, leading to a better understanding of pathogenic mechanisms occurring in patients where more than one specific protein may be aggregating in their neurons.

## Background

Amyotrophic lateral sclerosis (ALS) is an adult-onset progressive neurodegenerative disease characterised by the irreversible death of upper and lower motor neurons (Strong *et al*, 2005). This results the loss of voluntary muscle functions with death within 2-5 years after symptom onset. ALS exists under two different forms with most cases belonging to the sporadic ALS (sALS) group (90-95% of total cases) while just 10% of cases are related with inheritance elements following a predominantly autosomal dominant pattern (Masrori & Van Damme, 2020).

The neuropathological hallmark of ALS is the presence in motor neurons of neuronal cytoplasmic inclusions (NCIs) consisting of ubiquitinated protein aggregates. In 2006, it was determined that an RNA binding protein, Tar DNA binding protein 43 (TDP-43), represented the main component of these inclusions in the vast majority of ALS patients (Arai *et al*, 2006; Neumann *et al*, 2006). While this was confirmed in many subsequent studies, it is now also well described that several other proteins can also co-localise in these aggregates with varying abundance, as recently reviewed elsewhere (De Marchi *et al*, 2023). The importance of studying these co-aggregations is just beginning to emerge because of the possibility that they may influence both disease progression and duration (Blokhuis *et al*, 2013; Moda *et al*, 2023).

Among the proteins that have been observed to co-aggregate with TDP-43 inclusions, a potentially interesting co-aggregating factor is represented by Rho guanine nucleotide exchange factor (RGNEF) (Droppelmann *et al*, 2013; Keller *et al*, 2012). RGNEF is a 190kDa protein encoded by the *ARHGEF28* gene in humans that belongs to the GEF family and is acting as a survival factor in stress conditions (Cheung *et al*, 2017). Currently, it is the only protein described to be involved in ALS that combines a signal transduction activity through the GEF activation of RhoA together with the ability to bind and affect RNA processing (Miller *et al*, 2014). Under physiological conditions, RGNEF predominantly localises in the cytoplasm of motor neurons, with lower levels of localisation in the nucleus. Several functional domains have been described to be important for RGNEF GEF and RNA binding functions that include a Pleckstrin

Homology (PH) domain (Powis *et al*, 2023) and a Dbl Homology domain (Aghazadeh *et al*, 1998) that are present in the central part of the protein. The PH domain contains both a nuclear localisation signal (NLS) and a nuclear export signal (NES) that act independently to each other to determine RGNEF localisation in cells (Tavolieri *et al*, 2019). This domain is not important just for the localisation of the protein, but together with the Dbl homology domain is crucial also for the guanine exchange function activity of the protein (Droppelmann *et al*, 2014).

From a functional point of view, one of the best-known properties of RGNEF is in regulating the stability of neurofilament (*NF*) mRNA (*NEFL* mRNA). This could be important for ALS pathology because the cytoskeletal architecture organisation of the neurons is related to neurofilaments and one of the peculiar features of ALS is the progressive decrease *in NEFL* mRNA levels (Feneberg *et al*, 2018; Steinacker *et al*, 2016; Vacchiano *et al*, 2021).

Specifically, RGNEF has been reported to be able to bind *hNEFLs* mRNA 3’ untranslated region (UTR) resulting in the destabilisation of the transcript. Interestingly, TDP-43 acts as a stabilisation factor for *hNEFL*, reducing its degradation rate, thus suggesting a complementary functional role to that of RGNEF (Droppelmann *et al*., 2013).

To further address this matter, we have now characterized the wider mechanistic connection between these two RNA binding proteins and show that they possess a considerable overlap in regulating genes of the pathway related to axon guidance and synaptic functioning, whose disruption is a well-known effect caused by TDP-43 dysregulation in ALS patients (Lepine *et al*, 2022). From a mechanistic point of view, our experiment shows that the connection between these two proteins occur at the level of long-intron splicing, further supporting and extending the hypothesis that disruption of long-intron processing is a key functional aspect of TDP-43 pathology (Lagier-Tourenne *et al*, 2012). Our results highlight the complexity of protein aggregation in neurodegenerative processes and suggest that co-aggregating proteins in TDP-43 proteinopathies should be considered for a better understanding of disease onset and progression.

## Materials and Methods

### Cell growth and gene silencing in SH-SY5Y cells

Human neuroblastoma SH-SY5Y cell line (ECACC) was cultured up to 20 passages in Dulbecco’s modified Eagle’s medium (DMEM)/Nutrient Mixture F-12 Ham (Sigma-Aldrich), supplemented with 15% fetal bovine serum (FBS) (Sigma-Aldrich), 1% MEM Non-essential Amino Acid Solution (100X) (Sigma-Aldrich), and 1% Antibiotic-Antimycotic-stabilized suspension (Sigma-Aldrich) at 37°C with humidified atmosphere of 5% CO2.

SH-SY5Y cells underwent gene knock-down against two different genes: *TARDBP* and *ARHGEF28*. The sequence of the siRNA used to silence *TARDBP* was: 5’-gcaaagccaagaugagccu-3’, while the siRNA sequence to deplete *ARHGEF28* is the following: 5’-cgaccgaauuggagauauu-3’. Two different siRNA were used as negative controls. An siRNA against the firefly luciferase (siLUC) was used as a negative control for *TDP-43*: 5’-uaaggcuaugaagagauac-3’ while a silencer select negative control siRNA was used as a control for *RGNEF* (Catalog: 4390844, LifeTechnologies, ThermoFisher). Genes knock-down has been achieved through one round of silencing through Lipofectamine RNAiMAX (Invitrogen, ThermoFisher) according to the manufacturer’s instructions. The silencing of TDP-43 and RGNEF was performed by plating 8 x 10^5^ cells in p35 plates and using a reverse transfection with a mixture composed of: 150µL of Opti-MEM (LifeTechnologies), 3µL of 40µM siTDP-43 and control, or 10µM siRGNEF and control and 9µL of Lipofectamine RNAiMAX reagent (Invitrogen). The final concentration of the siRNAs in the plate was 80nM for siTDP-43. The reaction was performed at day 0 and was left for 48 hours. The final concentration in the plate for siRGNEF was 20nM and a first round of reverse transfection was performed in day 0, while a second round of forward transfection was performed in day 1 following the same protocol. Cells were collected on day 2 and prepared for western blot and and/or gene expression analysis.

### Sodium arsenite treatment

Cells were treated with 0.5 mM sodium arsenite for 40 minutes to induce cellular stress. After media substitution, cells were collected and prepared for Western blot and RNA extraction as previously described (Colombrita *et al*, 2009).

### Cycloheximide treatment

Cells were treated with cycloheximide (CHX, ThermoScientific) in a final concentration of 100µg/mL for 20 hours in dark conditions to induce the protein synthesis blockage. After media substitution, cells were collected and prepared for western blot and RNA extraction.

### Actinomycin D treatment

Cells, previously knocked-down for RGNEF and TDP-43, were treated with 5µg/mL actinomycin D (ActD, Sigma-Aldrich) and collected after 0h, 1h, 2h and 4h. Cells were then prepared for western blot and RNA extraction.

### Protein expression analysis

Cell pellets were resuspended in a lysis solution composed by 1 X Phosphate Saline Solution (PBS) supplemented with 1 X Complete Protease Inhibitor Cocktail (Roche) and then sonicated with a BioRuptor UCD-200 (Diagenode, Belgium) at high power. 30 µg of TDP-43 protein extract or 50µg of RGNEF extract was then resuspended in 1 X NuPAGE LDS Sample buffer (4 X) (Thermo Fisher Scientific) and denaturated at 85°C for 5 min. Western blots were performed using standard protocols using PVDF blotting membrane (Millipore, Merck). Membranes were then incubated with specific primary antibodies (**Suppl. Table 1**). Target luminescence was detected using Luminata Classico Western HRP substrate (Millipore, Merck) or Super Signal West Femto, Trial Kit (Thermo Fisher Scientific). Images were acquired with Alliance 9.7 Western Blot Imaging System (UVITEC).

### Real-time quantitative PCR analysis

RNA extraction was performed using RNeasy Mini Kit (Qiagen) with on column DNA digestion (Qiagen) following the manufacturer’s instructions. Reverse transcription was carried out at 37° C by random primers (Sigma-Aldrich) and Moloney murine leukemia virus (M-MLV) Reverse Transcriptase (Invitrogen). One microgram of total RNA was retrotranscribed and the cDNA was diluted 1:10 to the quantitative PCR (qPCR). Glyceraldehyde-3-phosphate dehydrogenase (GAPDH) and Ribosomal protein L30 (RPL30) housekeeping genes were used to normalise the results. The target/housekeeping gene sequences are listed in **Suppl. Table 2** and **Suppl. Table 3**. The quantification of gene expression levels was performed by quantitative Real-time PCR using PowerUp SYBR Green Master Mix (Applied biosystems) and QuantaStudio5 instrument (Applied biosystems). Results were analysed using QuantaStudio Design & Analysis software by ΔΔCt method. The mean of relative expression levels ± standard error of the mean (SEM) is reported in the corresponding graphs (*n*=3 independent experiments). Three technical replicates were considered for each sample. Nonparametric un-paired *t-test* was used as statistical test (GraphPad Prism software v8.0). Statistical significance was displayed as * *P* ˂ 0.05, ** *P* ˂ 0.01 and *** *P* ˂ 0.0001.

### RNA sequencing analysis of ARHGEF28 and TARDBP

mRNA library construction and RNA sequencing was performed by Novogene (https://en.novogene.com/) on three independent experiments obtained from SH-SY5Y cells depleted for ARHGEF28 (RGNEF) and TARDBP (TDP-43). RNA-seq analysis was performed using Illumina HiSeq 2500 instrument. CASAVA base recognition was used to transform the original raw data from Illumina to sequenced reads: 60 million paired-end 150 bp (PE150). Read trimming was performed in order to remove bases with low sequencing quality (meaning reads with more than 50% nucleotides quality less than five or reads with more than 10% reads uncertain nucleotides) and adapter sequences. For RNA sequencing of SH-SY5Y cells knockdown for ARHGEF28, trimmed reads were then mapped to the reference genome (GRCh37/hg19) using STAR software (v2.5). Differential gene expression analysis was carried out using DEseq2 R package (v1.22.2). On the other hand, RNA sequencing performed on SH-SY5Y depleted for TARDBP was already submitted to GEO database (GSE245303). In this work, trimmed reads were mapped to the reference genome GRCh37/hg19 and DEseq2 R package v.1.20.0 was used for differential expression analysis. For both datasets, the overall distribution of differentially expressed genes (DEGs) were evaluated using the following cut-off: up-regulated genes Fold Change (FC) >1.3 and *Padjust* < 0.05; down-regulated genes FC < 0.7 and *Padjust* < 0.05. Representation of RNA sequencing data was made by volcano plots using ggplot2 R package (v3.3.5). Over-represented KEGG (Kyoto Encyclopedia of Genes and Genomes) pathway analysis was also performed on the RNA sequencing data using GOseq R package (version 3.14.3).

### RNA-immunoprecipitation (RNA-IP) assay

SH-SY5Y cells were transfected with 16µg of His-tagged siRNA resistant RGNEF plasmid, with Flag-tagged siRNA resistant TDP-43 or pRc/CMV control vector (Invitrogen), following the manufacturer’s instructions. Cells were seeded in a 10mm tissue culture dish and grown under their normal conditions in order to reach 80% confluence on the day of transfection. 24 hours after the seeding, transfection was performed using a mixture composed by: 450 µL of EC buffer (Qiagen), 12 µL of Enhancer (Qiagen), 16 µg of plasmid and 15 µL of Effectene (Qiagen). Cells were collected after 48 hours and prepared for the RNA-IP experiment. RNA-IP was performed using Pierce^TM^ Classic Magnetic IP/Co-IP Kit (ThermoScientific), according to the manufacturer’s instructions. First an immune complex composed by the cell lysate and the specific antibody was prepared and let to react for 24 hours at +4°C. The reaction was performed using antibodies as following: 5 µL of mouse monoclonal anti-FLAGM2 (Sigma-Aldrich F1804), 5 µL of mouse polyclonal anti-HIS H8 to 6X His Tag (Abcam) or 5 µL of IgG from mouse serum as negative control. 24 hours later, the IP reaction was performed with the incubation for 1 hour of magnetic beads together with the immune complex. After the immunoprecipitation reaction, beads were washed three times and then the elution was performed by incubating the magnetic beads for 10 minutes with the Elution Buffer (ThermoScientific). The RNA was extracted using RNeasy Mini Kit (Qiagen) and an on column genomic DNA digestion was performed following the manufacturer’s instructions.

Reverse transcription of the RNA-IP samples and of 1% Input was carried out at 37° C by random primers (Sigma-Aldrich) and M-MLV Transcriptase (Invitrogen). cDNA was diluted 1:3 to perform RT-qPCR analysis. Quantification was performed using PowerUp SYBR Green Master Mix (Applied biosystems) and QuantaStudio 5 instrument (Applied biosystem). 1% input RNA fraction Ct value was used to normalise each RNA-IP fraction Ct value (IgG, anti-Flag and anti-HisTag). In each corresponding graph, the mean of the relative expression levels ± SEM (*n*= 3 independent experiments) is reported. Three technical replicates were considered for each sample. GAPDH was tested as nonspecific target. Multiple *t-test* was used as statistical test (GraphPad Prism software v8.0). Statistical significance was displayed as * *P* ˂ 0.05.

### Reverse transcription, RT-PCR

To detect the presence of cryptic exons, PCR primers were designed in the exons that were flanking the predicted region of cryptic exons inclusion. PCR reaction was prepared using gene-specific primers (20µM, Eurofins, primer sequences in **Suppl. Table 4**) and PrimeSTAR^®^ Max DNA Polymerase (Takara) following the manufacturer’s instructions.

The amplification was performed using MiniAmp Plus Thermal Cycler (Applied biosystem) following the DNA polymerase manufacturer’s instructions (30 amplification cycles). PCR products were separated by capillary electrophoresis Qiaxcel^®^ DNA High Resolution Kit (Qiagen).

### Immunofluorescence analysis

3 X 10^5^ SH-SY5Y cells were seeded and *ARHGEF28* and *TDP-43* knock-down was performed as previously described in the section Gene knock-down in SH-SY5Y cells. 24 hours after the second round of silencing, cells were washed once with 2mL of 1X PBS, fixed with 2mL of 3.2 % P-formaldehyde-PBS (Electron Microscopy Science) and incubated with primary antibodies diluted in 2% BSA/PBS (for a list see **Suppl. Table 5**). Images were acquired using Axioscope 5 microscope equipped with Axiocam 202 mono camera (Zeiss, Oberkochen, Germany), and a 63X objective. For the image analysis, ZEN 3.2 (blue edition) software (Zeiss, Oberkochen, Germany) was used.

## Results

### Effects of TDP-43 and RGNEF knockdown on their expression

Considering the critical role for TDP-43 in ALS pathogenesis, we first wondered if potential changes in TDP-43 expression could be observed after RGNEF depletion or, conversely, whether changes in TDP-43 could affect RGNEF expression. We then performed *RGNEF* and *TDP-43* knock-down on SH-SY5Y cells and we evaluated each silencing efficacy by WB (**Suppl. Fig. 1A and 1C**, respectively) and by RT-qPCR in the presence of siRGNEF and siTDP-43 **(Suppl. Fig. 1B and 1D**, respectively). Regarding RGNEF siRNA, no significant changes in TDP-43 expression were detected by WB and by RT-qPCR (**Fig. 1A**). In contrast, the analyses revealed a significant decrease in RGNEF protein and mRNA levels when TDP-43 was reduced (**Fig. 1B**). We then performed an RNA-IP assay to determine whether RGNEF could bind directly *TARDBP* mRNA or *vice versa*, laying the basis for potential explanations for this observation. As expected from the lack of expression changes, the RNA-IP assay did not show any significant binding between RGNEF protein and TDP-43 mRNA (**Fig. 1C**). On the other hand, the pulldown of *RGNEF* mRNA was enriched in Flag-tag TDP-43 transfected cells (**Fig. 1D**), suggesting that *RGNEF* mRNA could indeed be bound by TDP-43.

**Fig. 1:**
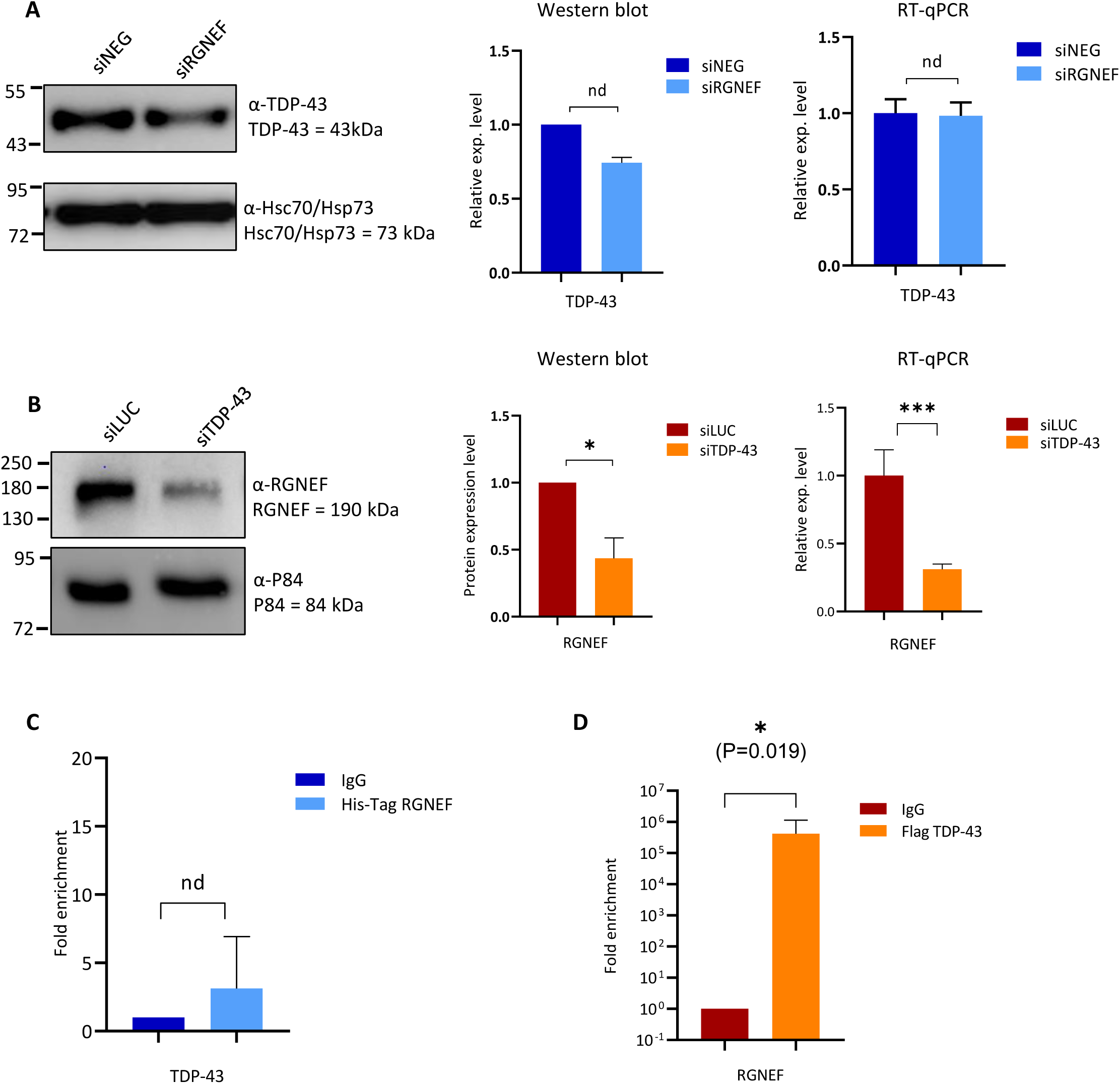
RGNEF effects over TDP-43 expression and TDP-43 effects over RGNEF expression. **(A)** Western blot analysis and relative quantification of TDP-43 protein level between controls and RGNEF depleted conditions (left side). On the right side of the panel the relative expression level of TDP-43 mRNA between controls and RGNEF depleted conditions is shown. siNEG: negative control; siRGNEF: samples silenced for RGNEF. **(B)** Western blot analysis and relative quantification of RGNEF protein level between controls and TDP-43 depleted conditions. siLUC: negative control; siTDP-43: samples silenced for TDP-43. **(C)** *TDP-43* mRNA enrichment after His-Tag RGNEF immunoprecipitation. The enrichment was evaluated by RT-qPCR; IgG: RNA immunoprecipitated negative control; His-Tag: RNA immunoprecipitation of target gene. Y axis values in logarithmic scale. **(D)** *RGNEF* enrichment after Flag TDP-43 immunoprecipitation. Enrichment evaluated by RT-qPCR as relative expression between the control and the Flag TDP-43 transfected samples; IgG: RNA immunoprecipitated negative control; Flag: RNA immunoprecipitation of target gene. Statistical analysis performed on three independent experiments. Each bar in the graphs reports the mean ± SEM. Nonparametric un-paired t-test was considered for statistical significance (* P˂ 0.05, ** P˂ 0.01, *** P˂ 0.001).

In parallel, we investigated the possible co-localisation between RGNEF and TDP-43 proteins in SH-SY5Y cells under basal conditions. As expected, TDP-43 was largely nuclear whilst RGNEF also displayed a significant nuclear localisation but seemed more cytoplasmic than TDP-43 **(Suppl. Fig. 2A)** as it is also demonstrated by the nuclear expression quantification of the two proteins **(Suppl. Fig. 2D)**. Interestingly, in addition to lower levels of RGNEF expression the TDP-43 knock-down induced a minimal shift of this protein to the cytoplasm (**Suppl. Fig. 2B and Fig. 2E**). Finally, RGNEF knock-down did not seem to alter TDP-43 localisation that remained predominantly in the nucleus (**Suppl. Fig. 2C and 2F**). Based on the reduction of RGNEF expression following *TDP-43* knock down, it was therefore considered very interesting to better characterize the global changes in transcriptome following reduction of RGNEF expression in the nucleus.

**Fig. 2:**
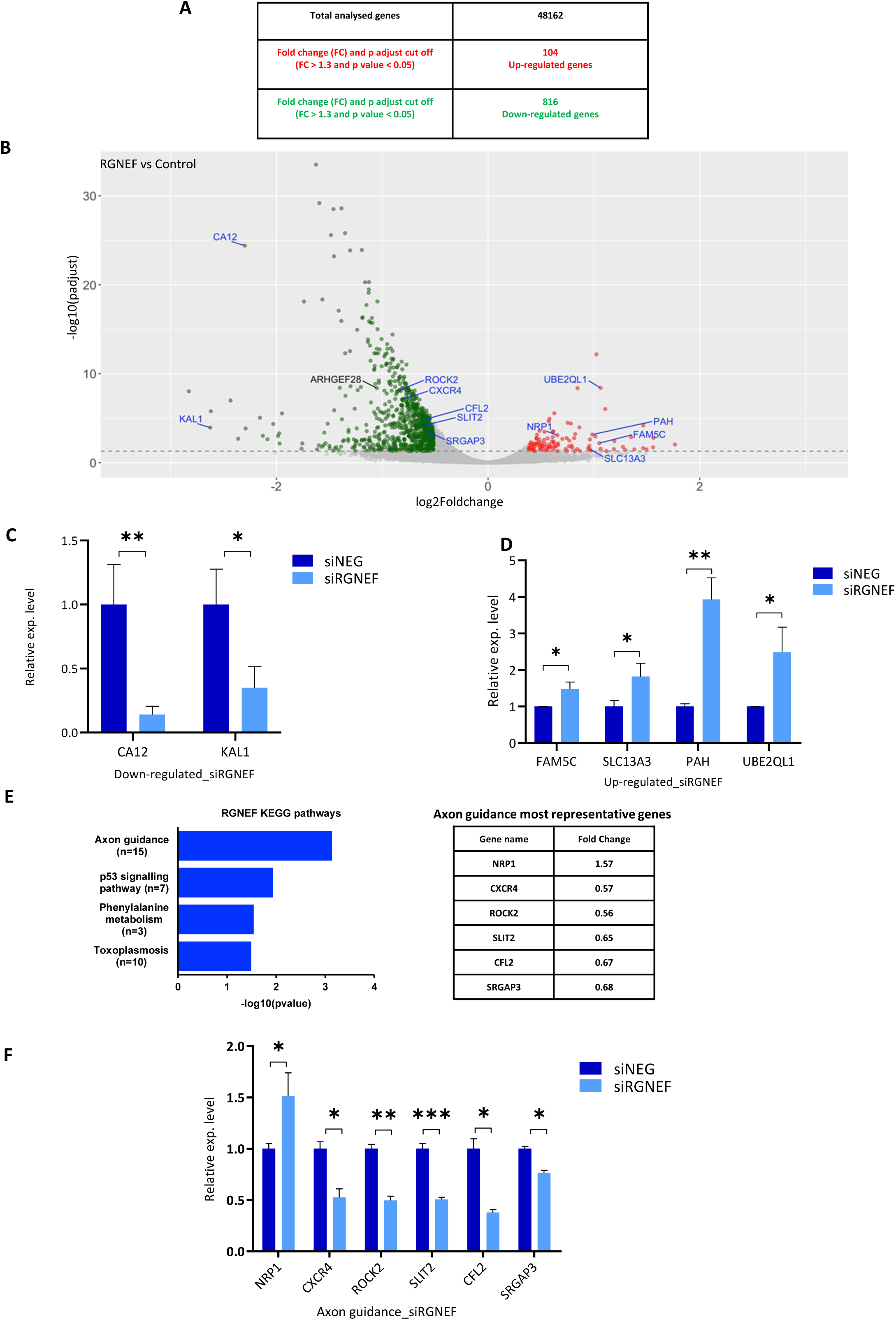
RNA-seq analysis of SH-SY5Y cells silenced for RGNEF. **(A)** Schematic table reporting the total number of analysed genes after RNA sequencing analysis. Red: up-regulated genes; green: down-regulated genes. (**B)** Volcano plot displaying the most significant differentially expressed genes in RGNEF*-*depleted samples compared to negative control. The down-regulated genes are shown in green while the upregulated ones are displayed in red. **(C)** Validation of relative expression levels of the top up-regulated targets among the top 10 up-regulated genes between controls and RGNEF depleted conditions. siNEG: negative control; siRGNEF: samples silenced for RGNEF. **(D)** Validation of relative expression levels of the top down-regulated targets between controls and RGNEF depleted conditions. **(E)** RGNEF KEGG analysis. List of pathways impacted by RGNEF depletion and differentially expressed genes among the axon guidance pathway. **(F)** Validation of relative expression levels of axon guidance related targets between controls and RGNEF depleted conditions. All the genes were selected basing on the Foldchange value choosing the ones with higher and lower values. Statistical analysis was performed on three independent RT-qPCR experiments. Each bar reports the mean ± SEM. Nonparametric un-paired t-test was considered for statistical significance (* P˂ 0.05, ** P˂ 0.01, *** P˂ 0.001).

### Identifying the RGNEF regulated targets in SH-SY5Y cells

To identify RGNEF-regulated genes, we performed RNAseq on samples deriving from *RGNEF* depleted SH-SY5Y cells. A negative control siRNA (siNEG) was used and three independent knock-down experiments were performed. From the RNA-seq analysis, out of 48,162 genes, the expression of 104 were up-regulated while 816 were down-regulated after siRNA treatment by applying a fold change (FC) cut-off of 30% compared to the control samples (**Fig. 2A**). To validate the RNA-seq data, the top 12 genes that showed significant differential expression after treatment with siRGNEF are shown in **Fig. 2B** as a Volcano plot. Among the differentially expressed genes (DEGs), *CA12* and *KAL1* were identified as the most affected genes following *RGNEF* knock-down and their down-regulation was validated comparing the relative quantification obtained via RT-qPCR analysis (**Fig. 2C**). Similarly, *FAM5A*, *SLC13A3*, *PAH* and *UBE2QL1* were also confirmed as the most up-regulated genes upon RGNEF knock-down (**Fig. 2D**). In parallel, KEGG analysis was performed to identify the most prominent pathway affected by RGNEF knockdown. In this analysis, 35 targets were identified as having significant gene enrichment (p≤0.05) and most of these targets were found to be associated with the axon guidance pathway (**Fig. 2E and Suppl. Table 6)**. Among these 35 genes, five down-regulated genes and the only up-regulated one were validated through RT-qPCR analysis, namely *CXCR4*, *ROCK2*, *SLIT2*, *CFL2*, *SRGAP3* and *NRP1* (**Fig. 2F**).

### RGNEF knock-down effects over ALS-associated protein expression and localisation

Because little is known about RGNEF possible influence on ALS-related genes expression, RT-qPCR analyses were performed to compare the relative expression levels of specific genes of interest between controls (siNEG) and samples silenced for *RGNEF* (siRGNEF) independently of the RNAseq data. First, we directed our attention on some the major neurodegeneration-associated genes (*FUS, Tau,* and *SOD1*) and on different hnRNPs that have been described to act as TDP-43 functional modifiers (Appocher *et al*, 2017; Cappelli *et al*, 2022). As shown in **Suppl. Fig. 3A**, the analysis revealed a modest but significant change in the gene expression levels of *hnRNPA0, hnRNPA2, hnRNPK, hnRNPU,* and *Tau* in siRGNEF treated samples. The same analysis was performed following sodium arsenite treatment to evaluate the effects of stress conditions and revealed a significant increase in the mRNA expression of *hnRNPA0* and *FUS* in RGNEF depleted samples compared to the control. However, unlike in normal conditions no changes were detected in *hnRNPA2*, *hnRNPK*, *hnRNPU* and *Tau* expression **(Suppl. Fig. 3B)**. For protein analysis, we directed our attention on the most significant genes revealed by RT-qPCR analyses, *FUS* and *hnRNPA0*. As expected from the RT-qPCR results, no significant change in FUS protein expression was detected under physiological conditions **(Suppl. Fig. 3C**). However, FUS levels were significantly up-regulated upon RGNEF knock-down under stress conditions **(Suppl. Fig. 3D)**, confirming the RT-qPCR analysis results. Likewise, hnRNPA0 protein levels were significantly higher upon RGNEF silencing only under stress conditions **(Suppl. Fig. 3F)** whilst they remained comparable between siRGNEF and controls in physiological conditions, despite the increase in mRNA levels **(Suppl. Fig. 3E)**. Considering the lack of changes in FUS protein expression under normal conditions, we then performed an immune-localisation analysis following RGNEF knockdown to evaluate an eventual shift in the nucleo-cytoplasmic localisation of this protein. However, FUS localisation remained mainly nuclear **(Suppl. Fig. 3G-H**). Taken together, these results suggest that RGNEF by itself does not have a high capacity to affect the expression of the most prominent ALS associated genes.

### TDP-43 and RGNEF common regulated targets

To identify common regulated mRNAs among RGNEF and TDP-43, the RNA-seq data of RGNEF depleted samples was merged with the dysregulated genes observed to occur in TDP-43 depleted SH-SY5Y cells. Out of 60448 total analysed genes, knockdown of *TDP-43* resulted in the upregulation of 291 genes and downregulation of 590 genes. Interestingly, among these 881 DEGs we found that the expression of 95 genes were altered following both RGNEF and TDP-43 silencing whilst 786 DEGs were TDP-43 specific and 825 were RGNEF specific (**Fig. 3A**). The top commonly regulated mRNAs were highlighted in the volcano plot corresponding to siRGNEF vs control treated cells (**Fig. 3B**) and were validated by RT-qPCR analysis. Interestingly, out of the most common up-regulated targets, many of these genes such as *GREM2*, *CGN*, *SLITRK6* showed an opposite change in expression when cells were treated with siRGNEF (**Fig. 3C**) compared to the siTDP-43 treated samples (**Fig. 3E**). In this validated list, *PTGER3* was the only exception that resulted being up regulated in both cases. Similarly, the same analysis was performed for the top down-regulated genes. In an analogous manner to the up-regulated genes, *SYPL1* and *CFL2* resulted down-regulated upon RGNEF depletion (**Fig. 3D**) but showed increased gene expression when TDP-43 was silenced (**Fig. 3F).** Nonetheless, exceptions were present. For example, *MPPED2* and *SRGAP3,* showed the same behaviour in RGNEF depleted cells and TDP-43 depleted ones, resulting in down-regulation in both TDP-43 and RGNEF silenced cells.

**Fig. 3:**
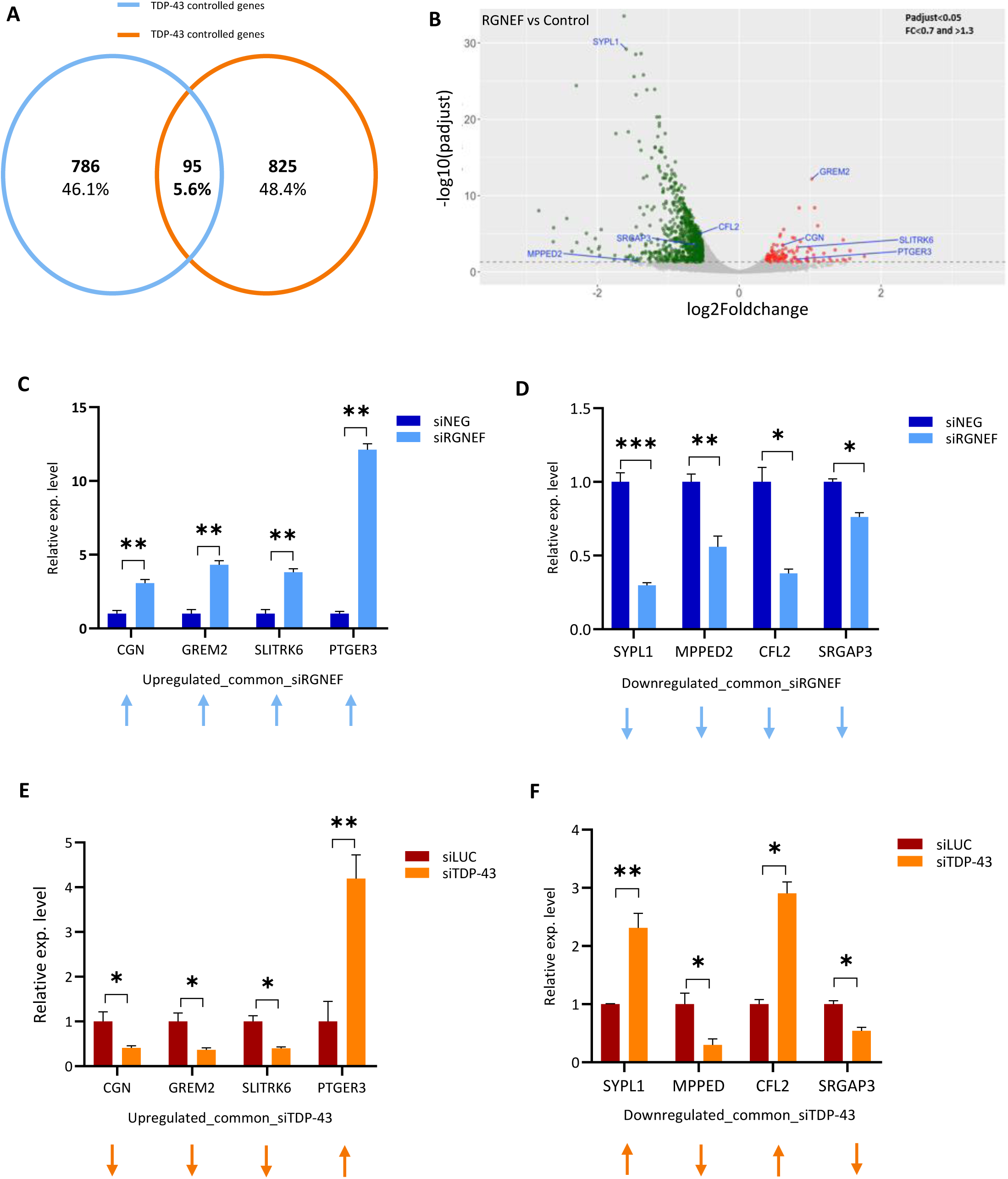
Bioinformatic analysis of common regulated genes between *RGNEF* and *TDP-*43. **(A)** Graph representing the percentage of gene regulated by TDP-43, by RGNEF and co-regulated. 95: n° of genes regulated by RGNEF and TDP-43 together; 786: n° of TDP-43 specific DEGs; 825: n° of RGNEF specific DEGs. **(B)** Volcano plot showing the common up- and down-regulated targets by *RGNEF* and *TDP-43* (n=95, the top genes are indicated). Green: down-regulated genes; red: up-regulated genes. **(C)** Validation of expression levels using RT-qPCR of up-regulated targets between controls and RGNEF depleted conditions. siNEG: negative control; si*RGNEF*: samples silenced for RGNEF. **(D)** Validation of expression levels using RT-qPCR of down-regulated targets between controls and RGNEF depleted conditions. **(E)** Validation of expression levels of up-regulated targets between controls and TDP-43 depleted conditions. siLUC: negative control; siTDP-43: samples silenced for TDP-43. **(F)** Validation of expression levels of down-regulated targets between controls and TDP-43 depleted conditions. For all validation studies statistical analysis was performed on three independent RT-qPCR experiments. Each bar reports the mean ± SEM. Nonparametric un-paired t-test was considered for statistical significance (* P˂ 0.05, ** P˂ 0.01, *** P˂ 0.001). Arrows indicating the regulation direction.

### Characterizing the effects of TDP-43 and RGNEF co-regulation on gene expression

A bioinformatics analysis of the gene length of the 95 co-regulated RGNEF and TDP-43 targets showed that compared to short (<15kb) (**Fig. 4A**) and middle genes (15-100 kb) (**Fig. 4B**), there was a significant enrichment in long genes in the co-regulated category (>100 kb) (**Fig.4C**). Specifically, whilst only 20.3 % of the total RGNEF regulated targets and 19.2% of genes regulated by TDP-43 were long genes, the percentage of long genes co-regulated by both proteins was increased to 27.4%. For this set of commonly regulated targets, we then wondered whether RGNEF and TDP-43 could directly bind these mRNAs. To achieve this, we performed an RNA-IP assay to identify the mRNAs bound by both TDP-43 and RGNEF. As shown in **Fig. 4D-E**, *SRGAP3, MPPED2, GREM2* and *CFL2* mRNAs were bound directly by both TDP-43 and RGNEF. In contrast, *SLITRK6* and *PTGER3* did not show any direct binding with these two proteins, and therefore their changes in expression probably originate from indirect effects. Transfection efficacy was evaluated by Western blot analysis (**Suppl. Fig. 4A-B**). Currently, there is no information about RGNEF binding sites, but an iCLIP analysis performed by Tollervey et al. (Tollervey *et al*, 2011) allowed us to perform a check for TDP-43 binding sequences.

**Fig. 4:**
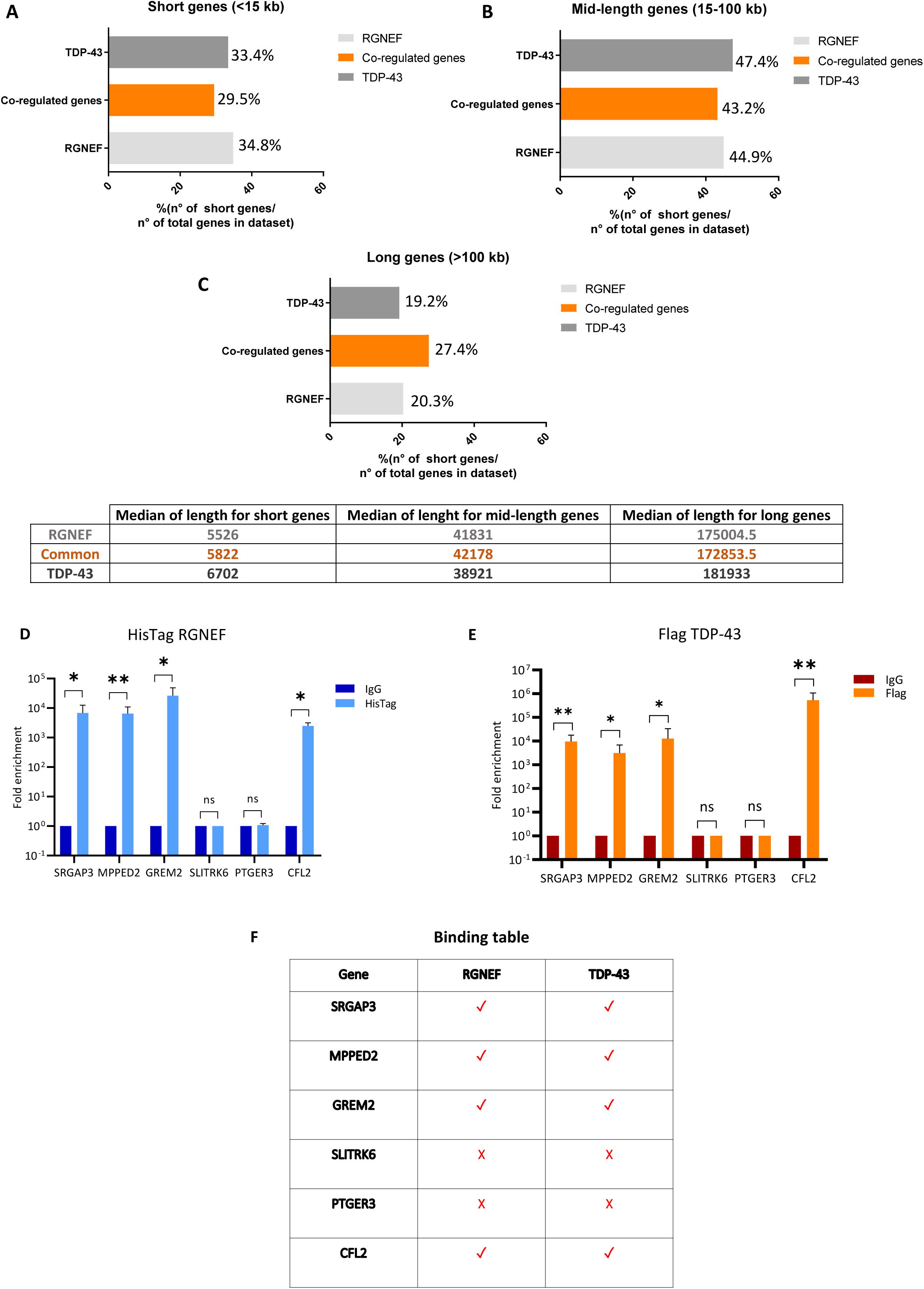
RNA-seq analysis of *RGNEF* and *TDP-43* common regulated genes and mRNA binding analysis. **(A)** Graph representing the percentage of RGNEF regulated genes, TDP-43 regulated genes and co-regulated genes among the short genes (˂ 15 kb). **(B)** Graph represented the percentage of RGNEF, TDP-43 and co-regulated genes among mid-length genes (15-100 kb). **(C)** Graph represented the percentage of RGNEF, TDP-43 and co-regulated genes among the long gene family (˃ 100 kb). Graphs and relative table showing the gene length analysis. **(D)** Genes directly bound by RGNEF using His-Tag RGNEF immunoprecipitation. IgG: RNA immunoprecipitated negative control; His-Tag: RNA immunoprecipitation of RGNEF gene. **(E)** Genes directly bound by TDP-43 using Flag TDP-43 immunoprecipitation. IgG: RNA immunoprecipitated negative control; Flag: RNA immunoprecipitation of TDP-43 gene. **(F)** Schematic table of the resulting mRNAs bound either by RGNEF and TDP-43. The pulldown enrichment was evaluated by RT-qPCR as relative expression between the control and the tag transfected samples; statistical analysis was performed on three independent RNA immunoprecipitation experiments. For *SLITRK6* and *PTGER3* no SD is reported because CT values were 45 for all the replicates. Y axis values in logarithmic scale.

Based on iCLIP, we observed that for *GREM2* and *MPPED2* multiple TDP-43 binding sites were located both on the exons but also in introns, whilst for *PTGER3* TDP-43 binding sites were just three and were all located at intronic level. Since it is also well known that TDP-43 preferentially binds TG/UG stretches (Buratti & Baralle, 2001) we also checked for the occurrence of these types of repeats in *GREM2*, *MPPED2* and *PTGER3* sequences via Ensembl Genome Browser (Ensembl Genome browser 110) and SnapGene (Dotmatics). As shown in **Suppl. Fig. 5A-C**, all these genes presented UG stretches at the level of intronic regions. Interestingly, in our experimental conditions *PTGER3* mRNA did not seem to be bound by TDP-43 or RGNEF according to RNA-IP, highlighting the concept that not all UG repeats in mRNAs can always be considered TDP-43 binders, as this could be impaired by RNA secondary structures or specific RNA modifications as recently described (McMillan *et al*, 2023).

### Differential long-intron processing rates of common regulated genes following TDP-43 and RGNEF depletion

Once the binding of RGNEF and TDP-43 to these mRNAs was confirmed, it was important to analyse which aspect of their mRNA processing steps could be altered upon knock-down of RGNEF and TDP-43.

As a result, we first considered the hypothesis that the binding with TDP-43 and RGNEF could affect RNA stability like it has been observed for many transcripts (Cappelli *et al*., 2022; Colombrita *et al*, 2012; Sidibe *et al*, 2021). However, treatment with ActD on TDP-43 and RGNEF-depleted cells did not show a significant change in *GREM2* mRNA stability following removal of these proteins **(Suppl. Fig. 6A-B**). For all these experiments, TDP-43 and RGNEF knock-down efficacy was evaluated by RT-qPCR **(Suppl. Fig. 6C-D**).

Secondly, we also wondered whether depletion of RGNEF or TDP-43 could induce the insertion of cryptic exons during the splicing process of these long-introns that might trigger alterations in their processing. Therefore, to investigate the possible presence of *GREM2* pseudo-exons, we amplified the mRNA sequence between exon 1 and exon 2 of this gene in the presence of CHX treatment to prevent nonsense mediated decay (NMD). However, no cryptic exon inclusion was detected in both siTDP-43 treated and not treated samples **(Suppl. Fig. 7A)**. In this experiment, CHX treatment effects were evaluated by western blot analysis showing a decrease in TDP-43 protein expression in samples treated with CHX; also silencing efficacy was evaluated by western blot analysis **(Suppl. Fig. 7B)**.

Finally, because many of these genes contain very long introns (>30 kbs), we wondered if the lack of RGNEF or TDP-43 could modify the speed long introns were processed from the RNA molecule. As shown in **Fig. 5A, left diagram** inspection of *GREM2* splicing profile from the RNAseq analysis revealed increased 118kb intron retention upon RGNEF between exon 1 and exon 2 (see **Suppl. Fig. 8A-B** for the whole set of RNAseq results). This increased intron retention was validated by RT-qPCR using primers that anneal to the exon and intron sequence that also showed up-regulation on the intron containing transcript when RGNEF was silenced but a down-regulation when TDP-43 was depleted (**Fig. 5A, right panels)**. These opposite results were in keeping with the observed effects of RGNEF and TDP-43 on gene expression where it was observed that *GREM2* mRNA expression was up-regulated following RGNEF knockdown (**Fig. 3C**) and down-regulated following TDP-43 knockdown (**Fig. 3E**). The same type of analysis was performed for *MPPED2* intron 4 that is 77kb long. However, in this case we observed a decrease in intron 4 containing pre-mRNA following both TDP-43 and RGNEF knockdown (**Fig. 5B, Suppl. Fig. 8C-D**). This result was in keeping with the observation that both TDP-43 and RGNEF downregulate the expression of this mRNA when they are silenced (**Fig. 3D-F**). Regarding another gene that contains long introns and whose expression is affected by TDP-43 and RGNEF knock-down in a positive manner we then analysed *PTGER3* that contains intron 2 which is 34kb long. For this gene, both the RNAseq and the RT-qPCR confirmed that knock down of RGNEF and TDP-43 up-regulated the amount of intron containing pre-mRNAs (**Fig. 5C, Suppl. Fig. 9A-B**). Again, this result was in keeping with the observation that both RGNEF and TDP-43 knockdown resulted in increased expression of PTGER3 mRNA (**Fig. 3C-E**). Finally, an effect on intron processivity was also observed for TDP-43 on the *RGNEF* pre-mRNA levels for the only long intron in its pre mRNA sequence that is represented by intron 3 which is 64kb long. Strikingly, it was observed that following TDP-43 knockdown the amount of pre-mRNA was decreased (**Fig. 5D, Suppl. Fig. 9C)**, in keeping with the effect of TDP-43 of down-regulating the expression of RGNEF following its removal (**Fig. 1B**).

**Fig. 5:**
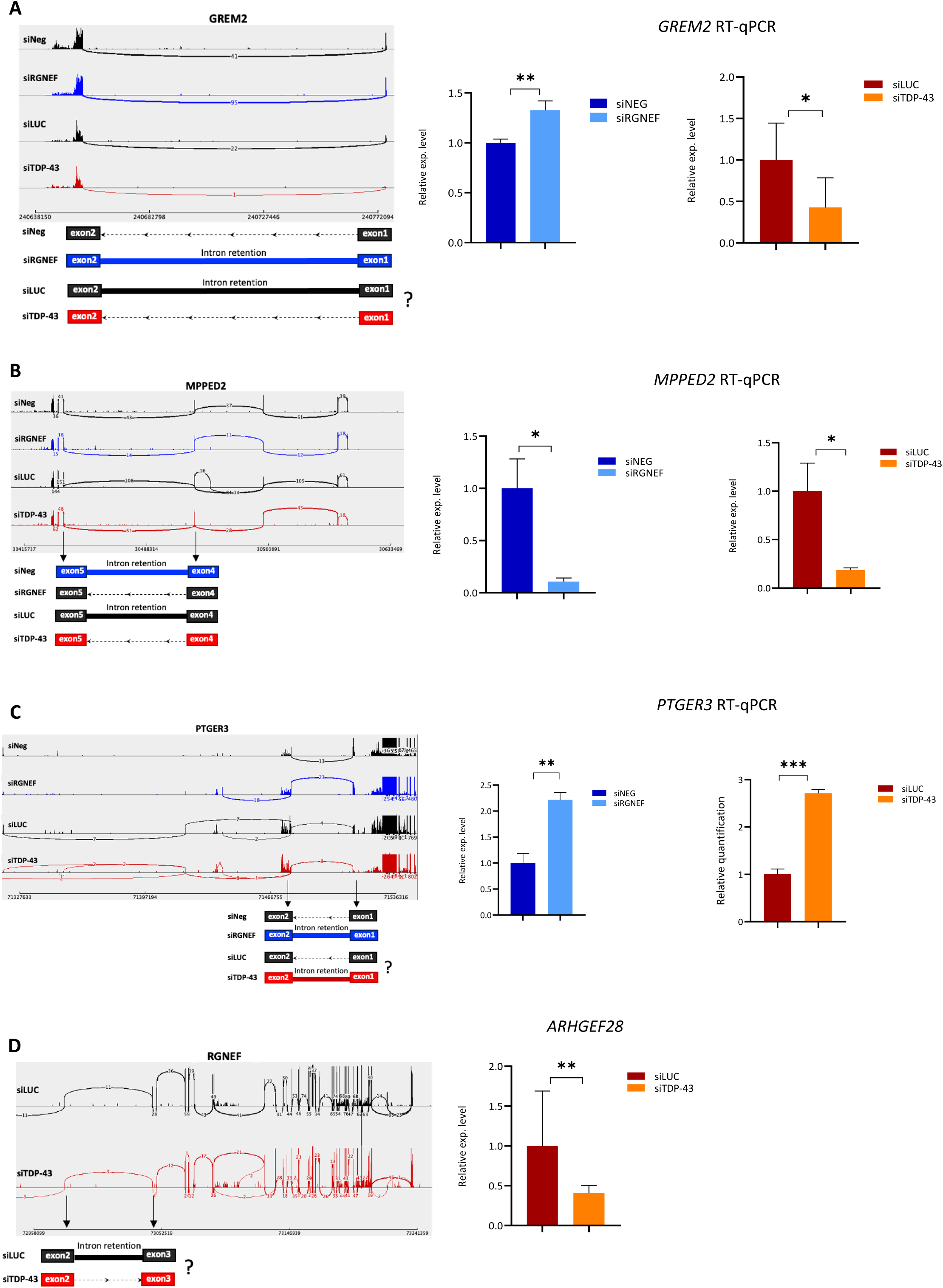
RGNEF and TDP-43 affect rate of long intron processing in common regulated genes. **(A)** On the left, sashimi plot of *GREM2* pre-mRNA was generated by IGV software using one exemplary RNA-seq sample obtained from SH-SY5Y cells depleted for siRGNEF vs control and siTDP-43 vs control. In each plot, exon regions are shown as “sashimi-like” regions, whereas intronic regions are shown as empty space between exonic regions. The number of junctions reads spanning from exons are reported as a “bridge” that crosses exons. A graphical representation of *GREM2* target exons and putative intron retention is also reported below the corresponding splicing profile. Relative expression levels of *GREM2* intron 1 levels between controls and depleted conditions were also reported on the right. *RGNEF* depleted conditions were showed in blueish, *TDP-43* depleted conditions were showed in reddish. siNEG: negative control; si*RGNEF*: samples silenced for *RGNEF*; siLUC: negative control; si*TDP-43* samples silenced for *TDP-43*. Statistical analysis was performed on three independent RT-qPCR experiments. Each bar reports the mean ± SEM. Nonparametric un-paired t-test was considered for statistical significance (* P˂ 0.05, ** P˂ 0.01, *** P˂ 0.001). Primer complementarity regions for *GREM2* intron 2 are shown by arrows over the graphs. **(B)** The same type of experimental set up for the *GREM2* pre-mRNA was applied to the *MPPED2* intron 4 pre-mRNA and for **(C)** the *PTGER3* intron 3 pre-mRNA whilst for the experiments in panel **(D)** the *ARHGEF28* (RGNEF) intron 2 pre-mRNA was analysed only for the siTDP-43 vs control conditions.

In conclusion, using several examples of genes whose expression is co-regulated by RGNEF and TDP-43 expression, our results suggest that the way these two proteins regulate gene expression (both in opposite or same ways) is by acting on the rate of long intron processivity.

## Discussion

TDP-43 and RGNEF have been described as proteins involved in ALS pathogenesis and both have been described as important actors on NCI formation, one of the hallmarks of disease. Although both proteins have been identified as RNA metabolism modulators, until today very little was known about their interaction apart from their ability to regulate in opposite direction the stability of *NEFL* RNA (Droppelmann *et al*., 2013). In this work, it was therefore interesting to see how much further the action of these two proteins could be overlapping beyond just the *NEFL* gene and what this could tell us about the possible consequences of a combined RGNEF and TDP-43 loss-of-function and we show that this overlap is part of a bigger picture well beyond the *NEFL* mRNA. More in general terms, the co-aggregation of different protein species in neurodegenerative diseases has become a topic of great importance since the observations that in patient’s brains, there is often aggregation of multiple proteins in addition to the primary pathology.

Specifically, our results show that the loss-of-function of RGNEF can mostly antagonize TDP-43 splicing for genes that are important for synaptic transmission. This is a particularly interesting finding because synaptic transmission is also one of the areas where there is considerable functional intersection between TDP-43 and another important protein involved in ALS pathology, FUS (Ling, 2018; Ratti & Buratti, 2016).

Interestingly, among the genes that we found to be regulated by RGNEF and are involved in the axon guidance, some of them have already been described in association with ALS pathology, such as *NRP1, CXCR4* and *ROCK2*. Specifically, Neuropilin 1 (NRP1) expression has been described as decreased in an ALS mouse model, specifically in the SOD1^G93A^ transgenic mice (Conti *et al*, 2014; Nijssen *et al*, 2018) and there is evidence that physiological Nrp1 is up-regulated in the central nervous system (CNS) and down-regulated in peripheral nervous system (PNS) after axonal injury or degeneration (Korner *et al*, 2019). Likewise, C-X-C chemokine receptor type 4 (*CXCR4*), also known as fusin, has been recently shown to play an important role in neuroinflammation and in neurogenesis (Yan *et al*, 2022) and its increased expression was also previously described in the ALS mouse model SOD1^G93A^ in which its expression was decreased (Luo *et al*, 2007). Moreovoer, its overexpression has also been reported to occur in human oligodendrocyte-like cells together with an enlargement of the axons of motor neurons in ALS (Andres-Benito *et al*, 2020; Yan *et al*., 2022).

Striking, CXCR4 was observed to decrease in microglia inflammatory markers and reduction in blood-brain-barrier permeability and this could lead to a better survival rate (Rabinovich-Nikitin *et al*, 2016; Yan *et al*., 2022). The same inhibition was also described as able to delay or prevent the ALS pathology progression (Janssens *et al*, 2018; Rabinovich-Nikitin *et al*., 2016; Yan *et al*., 2022). Finally, regarding *ROCK2*, the possible therapeutic potential of rho associated protein kinase (ROCK) inhibitors has been studied in different neurodegenerative disease models like spinal muscular atrophy (Bowerman *et al*, 2012), Huntington disease (Li *et al*, 2013) and Parkinson’s disease (Zhao *et al*, 2015). More attractive, ROCK2 inhibitors have been described as promising pharmacological molecules for ALS. In keeping with this view, a SOD1^G93A^ ALS mice model showed an increased levels of ROCK (Gunther *et al*, 2017; Takata *et al*, 2013) and the use of ROCK inhibitors pre-symptomatically seems to improve the motor functions and to increase the time of survival (Gunther *et al*., 2017; Roser *et al*, 2017). All these are in keeping with the previous observation that skeletal muscle biopsies of ALS patients showed increased levels of ROCK2 (Conti *et al*., 2014).

In addition to RGNEF-regulated genes, a neurological connection has also been described for some of the genes that are controlled by both RGNEF and TDP-43, such as *MPPED2* and *PTGER3*. More precisely *MPPED2* has been shown to play a role in human glioblastoma development, where the expression of *MPPED2* is significantly decreased as the severity of the glioblastoma increases and that its overexpression can slow down cell growth rate and inhibit their migratory ability (Pellecchia *et al*, 2021) Moreover, its deletion is associated with WAGR syndrome (Wilms tumour, aniridia, genitourinary abnormalities, range of development delays) in which also mental retardation has been reported (Schwartz *et al*, 1994; Tyagi *et al*, 2009). Besides *MPPED2*, *PTGER3* has also been described to be elevated in a rat model of Parkinson’ s disease (PD) (Schumann & Bar, 2022) and as one of the top ten hub genes in Alzheimer’ s disease, being able to interfere with the cyclic adenosine monophosphate (cAMP) formation whose alteration can be directly linked with disease (Maingret *et al*, 2017).

Taken together, the results of our study suggest that there is a complex interaction at the gene regulation level that will depend on whether a cell only has RGNEF aggregation, TDP-43 aggregation, or both. Our results suggest that depending on which scenario takes place, different genes will become mis-regulated or even compensated, with hard to predict results on the well-being of the neurons where this occurs (schematically depicted in **Fig.6A-C**).

**Fig. 6:**
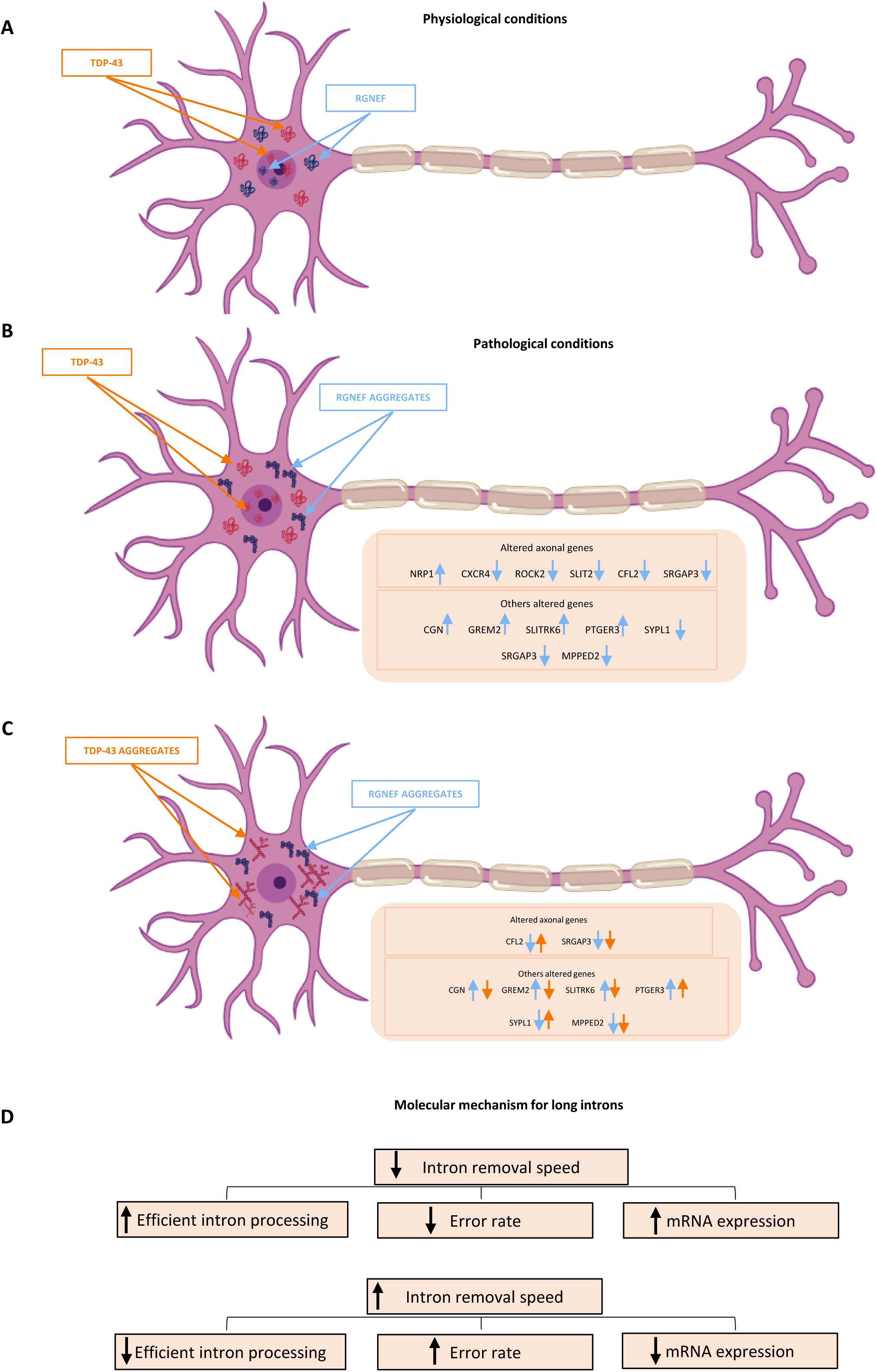
Possible consequences of long intron processing. **(A)** Schematic representation of TDP-43 and RGNEF normal physiological conditions in the cell. **(B)** Schematic representation of TDP-43 physiological and RGNEF pathological conditions in neurons with an indication of the major axonal genes being affected (the blue arrows indicate the direction of the change in the expression). **(C)** Schematic representation of what may occur to gene expression changes when both TDP-43 and RGNEF aggregate in neurons. The red arrows show the direction of expression change of each gene driven by TDP-43 absence. **(D)** Schematic representation of the possible consequences of fast vs. slow long intron processing mechanism by TDP-43 and RGNEF.

Most interestingly, at the mechanistic level we found that for the TDP-43/RGNEF co-regulated genes these two proteins seem to principally act on the rate of processing of long introns. The recognition and processing of long human introns where the correct 5’ and 3’ splice sites can be separated by thousands of Kbs (it has been estimated that >3000 human introns are longer than 50 kbp) has historically represented an enigma in understanding gene expression (Georgomanolis *et al*, 2016). To address this question, a few mechanisms have been put forward by looking at the way the splicing machinery can process long introns. The first was identified in *Drosophila* (Burnette *et al*, 2005; Duff *et al*, 2015) and in humans (Grellscheid & Smith, 2006), including human brains (Sibley *et al*, 2015) and has been called recursive splicing. In this process, long introns are removed in a stepwise manner by splicing at intronic sites that carry adjacent acceptor and donor splice sequences. More recently, this stepwise removal of introns has been confirmed in a single-cell, high-throughput approach where the rate of removal of thousands of introns in human bronchial epithelial cell can be measured in real time. Using this approach, the authors have shown that in many cases the loss of intron signal occurred before the RNA Polymerase II had reached the end of the signal, strongly suggesting that long intron removal proceeds in a stepwise fashion until the proper junction is reached (Wan *et al*, 2021).

Our work suggests that RGNEF and TDP-43 mostly act through their ability to increase or decrease long-intron removal speed and that this change in intron processivity would then explain the changes in mature mRNA expression.

Specifically, a higher intron removal speed could result in more recurrent errors in the splicing process, leading to an overall reduced gene expression as seen for the *MPPED2* gene both in RGNEF and TDP-43 depleted conditions or for *GREM2* when TDP-43 is silenced. Conversely, a reduced intron removal speed could result in a much more accurate splicing process, reducing the error rate and resulting in an overall higher gene expression (**Fig. 6D**). This would be the case for *GREM2* in RGNEF depleted conditions and of *PTGER3* in absence of both RGNEF and TDP-43.

In conclusion, our results support a very active role played by RGNEF in controlling key axon guidance genes and the view that its aggregation may be functionally important in ALS pathological processes. In addition, the overlap of RGNEF functions with TDP-43 and the observation that TDP-43 can control RGNEF expression levels is expected to add another layer of functional complexity to the possible consequences of TDP-43 aggregation in neurons. In fact, as previously shown for neuronal and muscle cell types (Susnjar *et al*, 2022), the number of genes altered by TDP-43 aggregation in cells is very sensitive to the local environment.

This situation of multiple protein aggregation is increasingly common in the ALS spectrum and makes it increasingly similar to what has been observed for FTD, where many hnRNP proteins have been recently describe to aggregate in specific neuronal types with and without TDP-43 pathology, such as hnRNP K (Bampton *et al*, 2021), hnRNPs R and Q (Gittings *et al*, 2019), and hnRNP E2 (Davidson *et al*, 2017) with many possible consequences on the well-being and survival of the neurons.

In conclusion, future lines of enquiry may include those aimed at understanding the specific vulnerability of specific neurons to both RGNEF and TDP-43 inclusions. In this respect, laser capture microdissection and single-cell RNA-seq approaches will prove helpful to address this issue. Finally, establishing whether post-translational modifications of RGNEF and TDP-43 are associated with their co-aggregation will also be insightful in this regard. In any case, our findings highlight the need to start considering the effect of multiple protein aggregation in neurodegenerative diseases.

## Acknowledgements

This work was supported by a generous donation from the Temerty Family Foundation to MJS and EB. MJS is supported by the Canadian Institutes of Health Research (CIHR). EB is supported by the Italian ALS Association AriSLA (project NOSRESCUEALS).

## Data Availability

Datasets discussed in this publication have been deposited in NCBI’s Gene Expression Omnibus and are accessible through the following GEO Series accession numbers: GSE248847 and GSE245303.

---

**Suppl. Fig. 1: RGNEF and TDP-43 knock-down efficacy in SH-SY5Y cells. (A)** Western blot analysis and relative quantification of RGNEF protein level between controls and RGNEF depleted conditions. **(B)** Relative expression level of *RGNEF* mRNA between controls and *RGNEF* depleted conditions. **(C)** Western Blot analysis and relative quantification of TDP-43 protein level between controls and TDP-43 depleted conditions. **(D)** Relative expression level of *TDP-43* mRNA between controls and *TDP-43* depleted conditions. Statistical analysis was performed on three independent experiments. Each bar in protein quantification reports the mean ± SD while bars of RT-qPCR quantification reports the mean ± SEM. Nonparametric un-paired t-test was considered for statistical significance (* P˂ 0.05, ** P˂ 0.01, *** P˂ 0.001). siNEG: negative control; siRGNEF: samples silenced for RGNEF. siLUC: negative control; siTDP-43: samples silenced for TDP-43.

**Suppl. Fig. 2: RGNEF and TDP-43 immuno-localisation. (A)** Immuno-localisation analysis of RGNEF and TDP-43 in normal conditions. DAPI: nuclear marker; RGNEF: α-goat Alexa Fluor 488; TDP-43: α-rabbit Alexa Fluor 594. **(B)** Immuno-localisation analysis of RGNEF under TDP-43 depleted conditions; DAPI: nuclear marker; TDP-43: α-rabbit Alexa Fluor 594. siLUC: negative control; siTDP-43 samples silenced for TDP-43. **(C)** Immuno-localisation analysis of TDP-43 under RGNEF depleted conditions. DAPI: nuclear marker; RGNEF: anti-goat Alexa Fluor 48. siNEG: negative control; siRGNEF: samples silenced for RGNEF. **(D)** Statistical quantification of RGNEF and TDP-43 expression levels in the nucleus and in the cytoplasm compared to the DAPI control. **(E)** Statistical quantification of RGNEF expression levels in the nucleus and cytoplasm of TDP-43 depleted samples compared to the siLUC control samples. **(F)** Statistical quantification of TDP-43 expression levels in the nucleus and cytoplasm of RGNEF depleted samples compared to the siNEG control samples. Each bar represents the mean ± SD. Nonparametric un-paired t-test was considered for statistical significance.

**Suppl. Fig. 3: RGNEF influence over neurodegenerative associated proteins. (A)** Relative expression levels of neurodegeneration-associated proteins and hnRNPs mRNA between controls and RGNEF depleted condition. **(B)** Relative expression levels of neurodegeneration-associated proteins and hnRNPS mRNA between controls and RGNEF depleted conditions in sodium arsenite treated samples. **(C)** Western blot analysis and relative quantification of FUS between controls and RGNEF depleted conditions. **(D)** Western blot analysis and relative quantification of FUS between controls and RGNEF depleted conditions of sodium arsenite treated samples. **(E)** Western blot analysis and relative quantification of hnRNP A0 between controls and RGNEF depleted conditions. **(F)** Western blot analysis and relative quantification of hnRNP A0 expression between controls and RGNEF depleted conditions of sodium arsenite treated samples. **(G)** Immuno-localisation analysis of FUS in RGNEF depleted conditions. DAPI: nuclear marker; FUS: α-rabbit Alexa Fluor 488. **(H)** Statistical quantification of TDP-43 expression levels in the nucleus of RGNEF depleted samples compared to the siNEG control samples. Statistical analysis was performed on three independent experiments. Each bar in protein and immune-locaisation quantification reports the mean ± SD while bars of RT-qPCR quantification report the mean ± SEM. Nonparametric un-paired t-test was considered for statistical significance (* P˂ 0.05, ** P˂ 0.01, *** P˂ 0.001). siNEG: negative control; siRGNEF: samples silenced for RGNEF; siNEG_ARS: negative control stressed with sodium arsenite treatment; siRGNEF_ARS: samples silenced for RGNEF and stressed with sodium arsenite treatment.

**Suppl. Fig. 4: HisTag RGNEF and Flag TDP-43 transfection efficacy. (A)** Western blot analysis of His-Tag RGNEF protein level between controls and RGNEF transfected conditions. pCMV: negative control; pHis-Tag RGNEF: samples transfected with His-Tag RGNEF. **(B)** Western blot analysis of Flag TDP-43 protein level between controls and Flag TDP-43 transfected conditions. pCMV: negative control; pFlag TDP-43: samples transfected with Flag TDP-43.

**Suppl. Fig. 5: iCLIP analyses and TG repeats presence in target genes. (A)** iCLIP analysis showing TDP-43 binding sites on *GREM2* mRNA. Sequence detail showing the presence of a TG stretch in its intron 1. **(B)** iCLIP analysis showing TDP-43 binding sites on *MPPED2* mRNA. Sequence detail showing the presence of a TG stretch in its intron 4. **(C)** iCLIP analysis showing TDP-43 binding sites on *PTGER3* mRNA. Sequence detail showing the presence of a TG stretch in its intron 1. (Adapted from Tollervey et al. 2011, Ensembl.org).

**Suppl. Fig. 6: mRNA stability assay**. **(A)** Actinomycin D assay to investigate the mRNA stability of *GREM2* transcript after TDP-43 silencing treatment. Relative RNA expression was evaluated by RT-qPCR at 0, 1, 2, and 4 hours after the treatment. Statistical analysis performed on three independent RT-qPCR experiments. Each point reports the mean ± SEM. Nonparametric un-paired t-test was considered for statistical significance. **(B)** The same type of experimental set-up was done to investigate the mRNA stability of *GREM2* transcript after RGNEF knock-down. **(C)** Relative *TARDBP* expression was evaluated by RT-qPCR to check the quality of TDP-43 silencing. **(D)** Relative *ARHGEF28* expression was evaluated by RT-qPCR to check the quality of RGNEF knock-down. Statistical analysis performed on three independent RT-qPCR experiments. Each bar reports the mean ± SEM. Nonparametric un-paired t-test was considered for statistical significance (* P˂ 0.05, ** P˂ 0.01, *** P˂ 0.001).

**Suppl. Fig. 7: Cryptic exon inclusion. (A)** Capillary electrophoresis analysis of *GREM2* expression between control and TDP-43 depleted conditions before and after cycloheximide treatment. CHX -: samples not treated with cycloheximide; CHX +: samples treated with cycloheximide. Primers complementarity regions in the *GREM2* pre-mRNA are shown on the left. **(B)** Western blot analysis of TDP-43 between controls and TDP-43 depleted conditions of CHX treated samples.

**Suppl. Fig. 8: Intron retention profiling of *GREM2* and *MPPED2* genes.** Sashimi plots was generated from the Integrative Genomics Viewer of RNAseq data obtained from SH-SY5Y cells depleted for RGNEF or for TDP-43 vs control. In each plot, exon regions are showed as “sashimi-like” regions, whereas intronic regions are showed as empty space between exonic regions. The number of junction’s reads spanning from exons are reported as a “bridge” that crosses exons. A graphical representation of exons and putative intron retention is also reported below the corresponding splicing profile. **(A)** Intron retention of *GREM2* in RGNEF depleted cells. **(B)** Intron retention of *GREM2* in TDP-43 depleted cells. The number of junction reads were not sufficient to detect intron retention. **(C)** Intron retention profile of *MPPED2* in RGNEF depleted cells **(D)** Intron retention profile of *MPPED2* in TDP-43 depleted cells. The number of junction reads were not sufficient to detect intron retention.

**Suppl. Fig. 10: Intron retention profiling of *PTGER3* and *ARHGEF28* genes.** Sashimi plots was generated from the Integrative Genomics Viewer of RNAseq data obtained from SH-SY5Y cells depleted for RGNEF or for TDP-43 vs control. In each plot, exon regions are showed as “sashimi-like” regions, whereas intronic regions are showed as empty space between exonic regions. The number of junction’s reads spanning from exons are reported as a “bridge” that crosses exons. A graphical representation of exons and putative intron retention is also reported below the corresponding splicing profile. **(A)** Intron retention profile of *PTGER3* in RGNEF depleted cells. **(B)** Intron retention profile of *PTGER3* in TDP-43 depleted cells. The number of junction reads were not sufficient to detect intron retention. **(C)** Intron retention profile of *ARHGEF28* in TDP-43 depleted cells. The number of junction reads were not sufficient to detect intron retention.

## SUPPLEMENTARY MATERIAL

**Supplementary Table 1.** Specific antibodies for western blot.

| Target protein | Manufacturer | Host | Dilution used | Dilution medium |
| --- | --- | --- | --- | --- |
| RGNEF | Abcam | Rabbit | 1:250 | 2%Milk/PBS-Tween |
| TDP-43 | Proteintech | Rabbit | 1:1000 | 2%Milk/PBS-Tween |
| Flag M2 | Sigma-Aldrich | Mouse | 1:1000 | 2%Milk/PBS-Tween |
| Myc | Abcam | Mouse | 1:1000 | 2%Milk/PBS-Tween |
| Tubulin | Home made | Mouse | 1:1000 | 2%Milk/PBS-Tween |
| hnRNP A0 | SigmaAldrich | Rabbit | 1:500 | 2% Milk/PBS-Tween |
| FUS | Abcam | Rabbit | 1:500 | 2% Milk/PBS-Tween |
| Hsc70/Hsp73 | Enzo Life Sciences | Rat | 1:3000 | 5%Milk/PBS-Tween |
| p84 | Abcam | Mouse | 1:1000 | 2%Milk/PBS-Tween |
| Anti-rabbit | Dako | Goat | 1:2000 | 2%Milk/PBS-Tween |
| Anti-mouse | Dako | Goat | 1:2000 | 2%Milk/PBS-Tween |
| Anti-rat | Bethyl | Goat | 1:2000 | 0.1%PBS-Tween |

**Supplementary Table 2.**
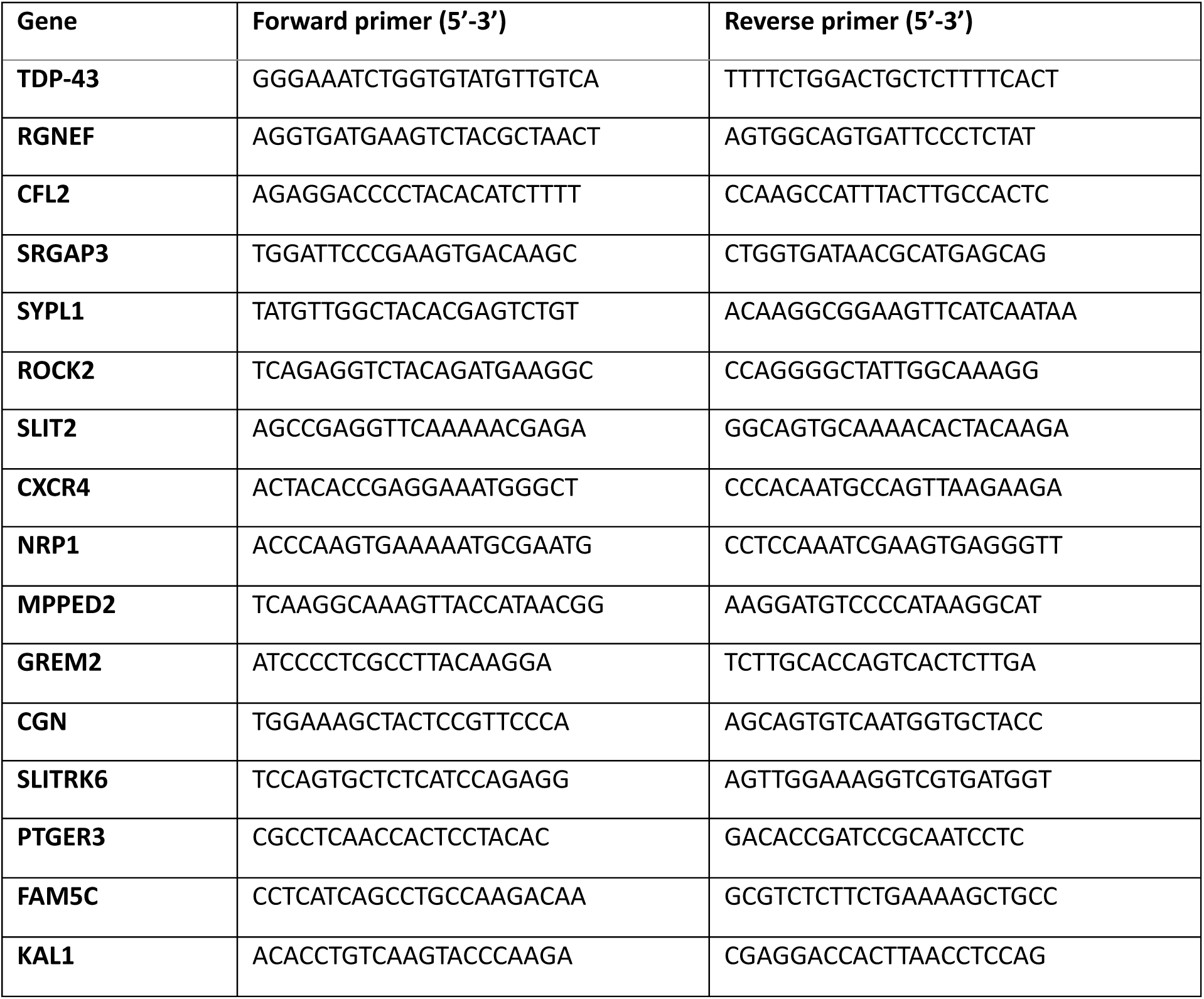
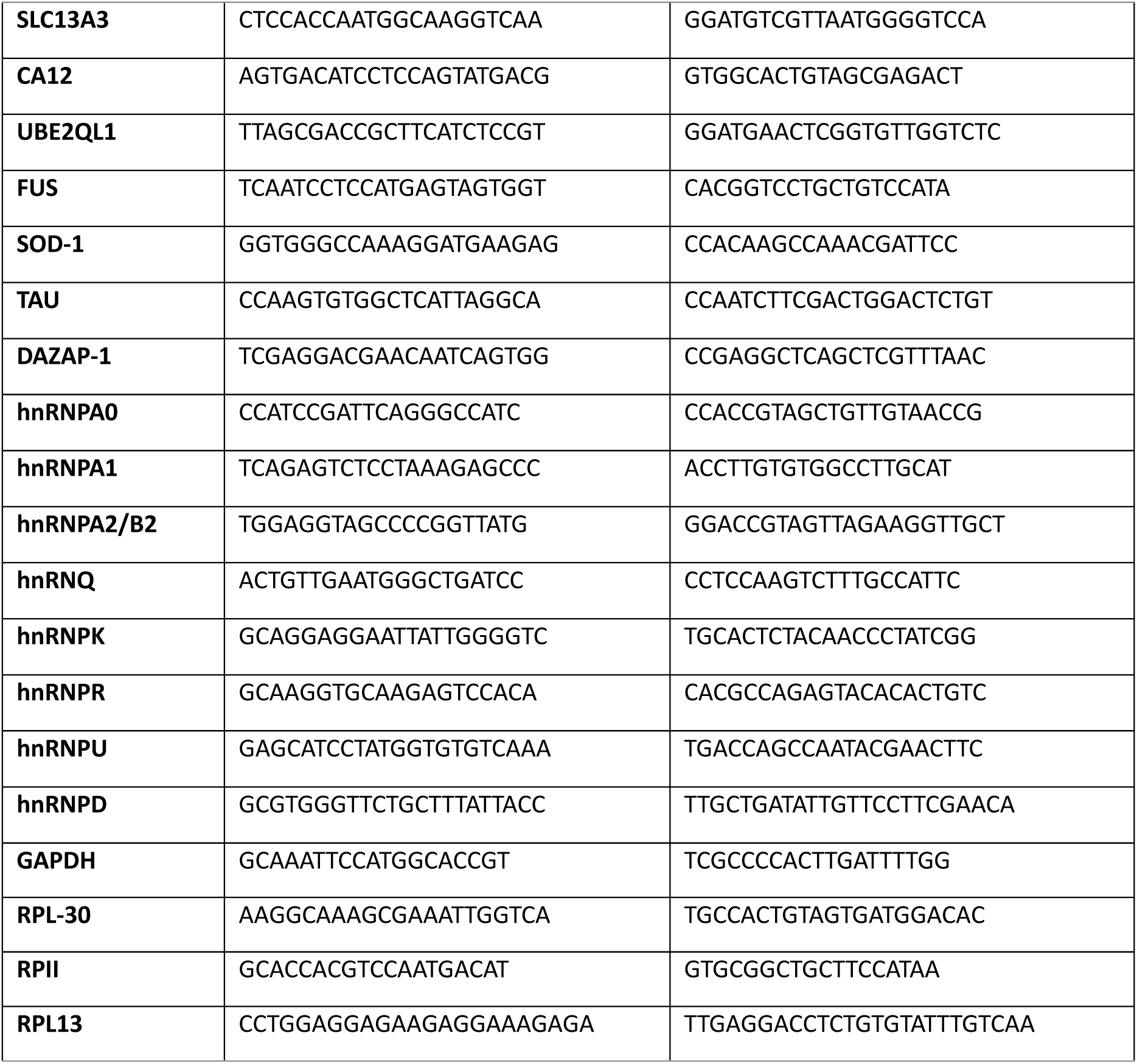
RT-qPCR primers.

**Supplementary Table 3.**
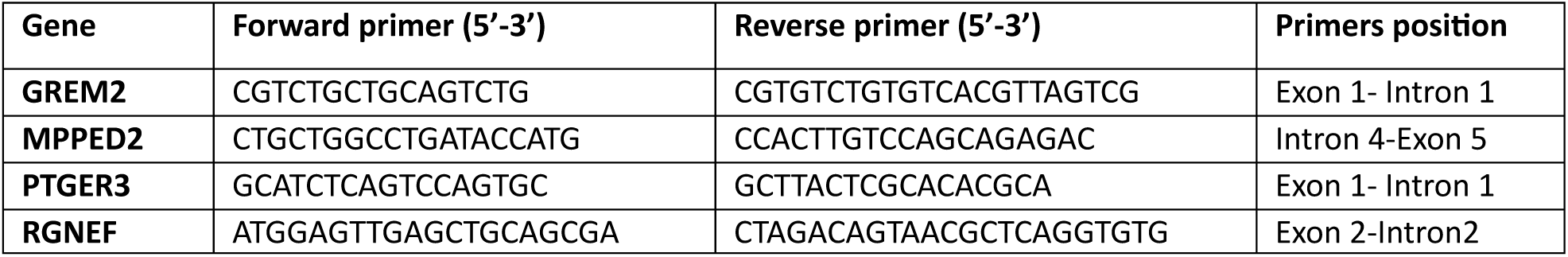
Primers for long intron processing RT-qPCR.

**Supplementary Table 4.**
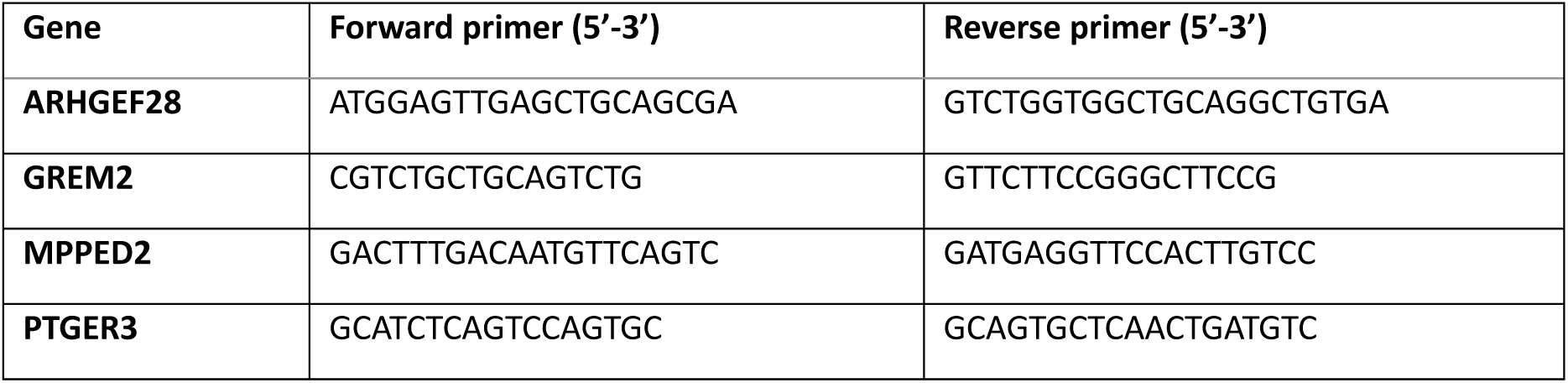
RT-PCR primers.

**Supplementary Table 5.**
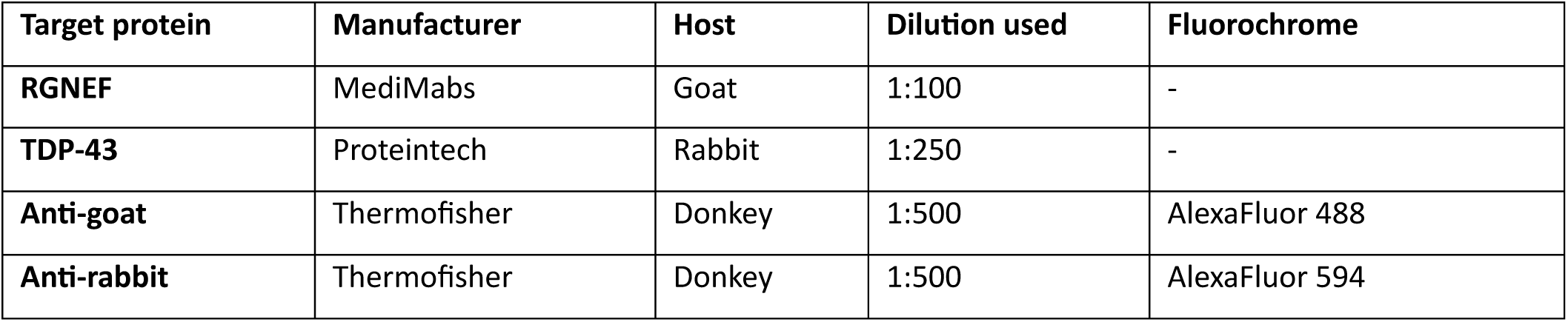
Immunofluorescence antibodies.

| Target protein | Manufacturer | Host | Dilution used | Fluorochrome |
| --- | --- | --- | --- | --- |
| RGNEF | MediMabs | Goat | 1:100 | - |
| TDP-43 | Proteintech | Rabbit | 1:250 | - |
| Anti-goat | Thermofisher | Donkey | 1:500 | AlexaFluor 488 |
| Anti-rabbit | Thermofisher | Donkey | 1:500 | AlexaFluor 594 |

**Supplementary Table 6.**
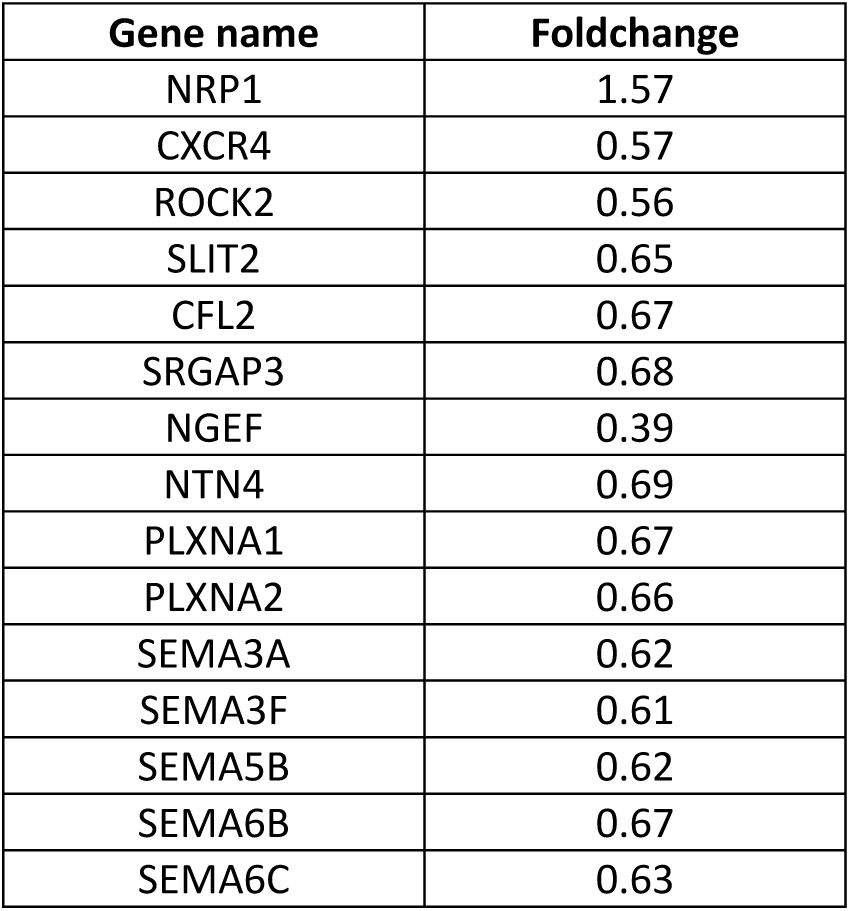
Representative table of KEGG analysis data of genes involved in axon guidance.

